# A New Ultradian Rhythm Linked to Protein Degradation and Synthesis in Mammalian Cells

**DOI:** 10.1101/2019.12.20.882811

**Authors:** L. Ghenim, C. Allier, P. Obeid, L. Hervé, J-Y. Fortin, M. Balakirev, X. Gidrol

**Author notes:** Correspondence to and.fr.

## Abstract

We describe a new ultradian rhythm that occurs during the interphase of the cell cycle in a wide range of mammalian cells, including both primary and transformed cells. The rhythm was detected by holographic lens-free microscopy that follows the individual histories of the dry mass of thousands of live cells simultaneously, each at a resolution of five minutes. Importantly, the rhythm was observed in inherently heterogeneous cell populations, thus eliminating synchronization and labeling bias. The rhythm is independent of circadian rhythm, has a period of 4 hours and is temperature-compensated. We demonstrated that the 4 hr rhythm is suppressed by proteostasis disruptors and is detected only in proliferating cells, suggesting that it represents the periodic dynamics of protein mass in growing cells.

**Brief teaser:** We have revealed a 4 hr rhythm in cell dry mass dynamics that seems to be general in proliferating mammalian cells.

## Introduction

Spatiotemporal order in a living system is believed to be a result of nonlinear chemical reactions and high variability. This leads to the generation of periodic oscillations that are characterized by a central timekeeping mechanism, which is defined as a clock for biological processes *(1)*. These rhythms allow organisms to adaptively cope with changing environments, such as the circadian clock imposed by Earth, thus conferring robustness and a competitive advantage. Many periodic biological processes have been described such as circadian rhythms, ultradian rhythms (periods shorter than a day) and the cell cycle. Some of the processes are endogenous, whereas others may be driven by external cues at the level of the organism. Nevertheless, the basic pacemaker of biological clocks appears to reside on a cellular level and is integrated into molecular networks of the cell. Studies in cultured cells have significantly advanced our understanding of the molecular mechanisms of the cell cycle and, more recently, of other cell-autonomous biological clocks. To detect periodicity in cell behavior, the expression of specific genes is usually measured by biochemical sampling or by using specific luminescent markers. Because the individual rhythms in a cell population are out of phase, they need to be synchronized by using an external stimulus. As a result, the most common approaches to studying biological clocks have a poor time resolution and may suffer from artifacts introduced by labeling or synchronization. On the other hand, our ability to study single-cell dynamics in asynchronous culture has been limited by the availability of quantitative phase imaging techniques for simultaneous, time-resolved, single-cell data acquisition from thousands of cells in parallel. Large population sampling is required to overcome the complex “*noisy”* behavior of a single cell and determine the standard characteristics of cell dynamics.

Recently, we have described a lens-free microscopy technique that allows for real-time measurements of dry mass with a precision of approximately 35 pg *(2)*. It is based on holographic interferences between the optical path that are propagated through a pinhole and waves scattered by the cell. The dry mass of the cell is calculated by integrating the phase of the diffracted light over the whole area of the cell *(2)*. Compared to conventional optical methods, lens-free microscopy provides a unique way to track thousands of live cells in real time and with a large field of view (35 mm^2^) without any labeling or synchronization. In the present work, we use this technique to demonstrate periodic oscillations in the cell dry mass. In the raw signal these are superimposed on a linear background that reflects cell growth.

## Results

### The 4 hr Rhythm

The experimental approach is illustrated in Fig. 1A. We used lens-free microscopy to analyze a large number of cells over several days by taking holographic images every 5 minutes. The reconstructed phase signal for each individual cell was used for cell segmentation and calculation of the cell dry mass *(2)* (see Fig. S1 to S4 and Materials and Methods for details). The data were then assembled to build individual cell trajectories and to reconstruct the histories of the cell dry mass. Mitotic events were readily identified from the corresponding abrupt changes in the cell dry mass concomitant with the appearance of a new cell trajectory (Fig. S3). We could then determine the cell cycle duration from the mean generation time between such events.

**Fig. 1.**
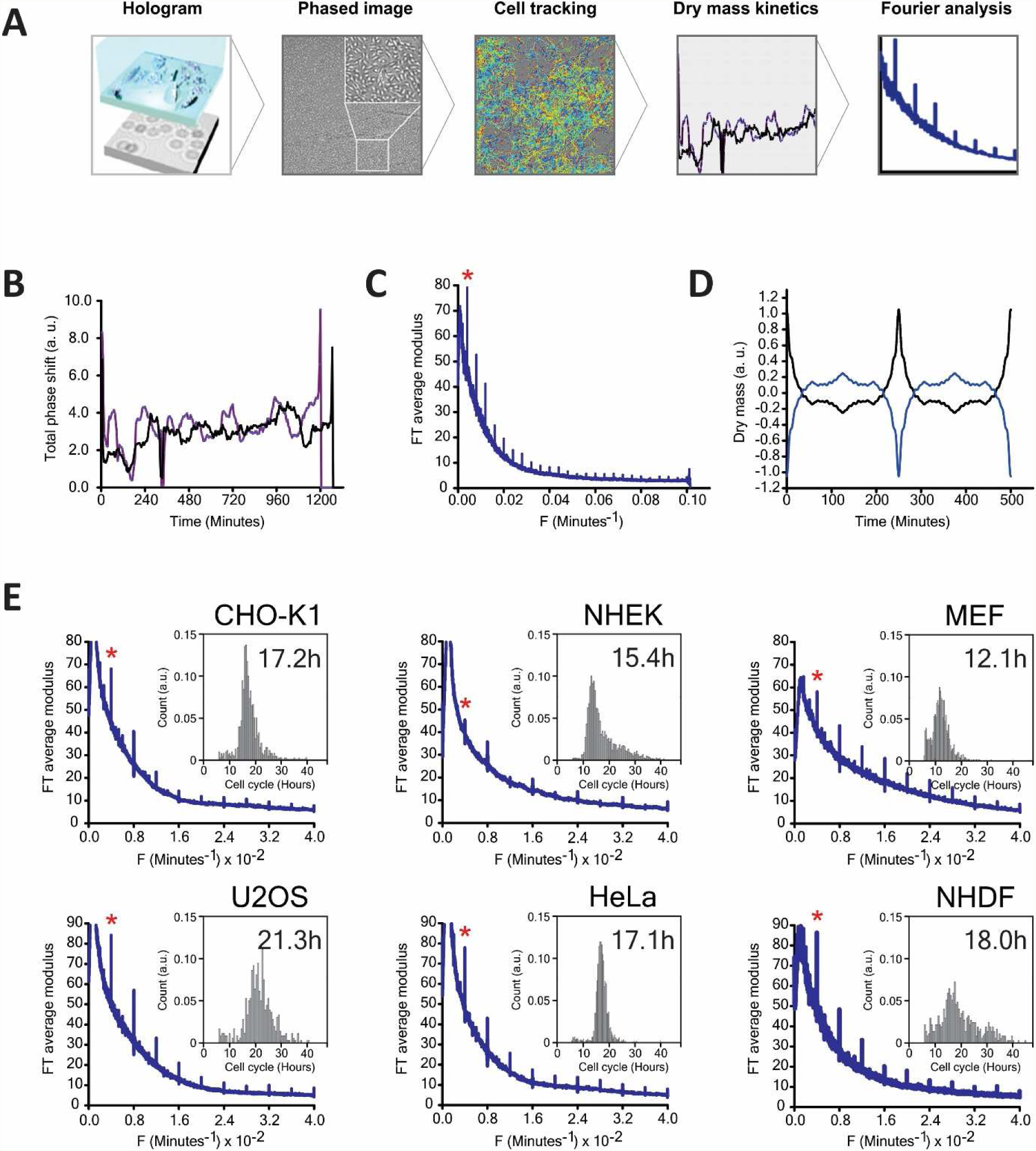
(A) A schematic view of the analysis of single-cell dry mass oscillations. (B) Individual histories of the dry mass of two individual mouse embryonic fibroblasts cells are shown during the interphase of the cell cycle. (C) The Fourier amplitude of the dry mass traces as a function of frequency averaged over many (N=702) MEF cells. We observe a dominant frequency (marked with an asterisk) of 0.004 min^-1^ (4.1 hours) as well as higher harmonics superimposed on a background of 1/f noise. (D) Reconstruction of the corresponding signal of dry mass by means of an inverse Fourier transform (with two possible solutions). (E) The 4 hr rhythm is observed in different cell lines. In the insert, the distribution of the cell cycle times is shown, and the mean value is in hours.

Cell dry mass dynamics were analyzed during the interphase of the cell cycle starting from the first detected mitotic event until the second (following) mitotic event or until the track was lost. For each cell j, after removing the linear background, we computed the Fourier transform (FT) of the dry mass change over the duration of the interphase trajectory T(j). The averaged absolute amplitude was properly normalized to take into account the different lengths of the traces (see Materials and Methods for details).

Analysis of the dry mass history in mouse embryonic fibroblasts (MEFs) revealed a dominant frequency of 0.004 min^-1^, along with its higher harmonics, superimposed on a background of 1/f noise (Fig. 1B). Excluding the 1/f noise, reconstruction of the raw signal by applying an inverse Fourier transform suggests that the cell dry mass shows periodic symmetric spikes every 4.0 hours. The signals attain the maximum amplitude and return to the background value with a characteristic time of approximately 30 min (Fig. 1B). Note that in the reconstructed function, the average amplitude for the change in the dry mass (the strength of the oscillator) depended on the amplitude of the principal FT component, whereas the form of the signal was defined by all harmonics.

Using this procedure, we studied several cell lines: Hela, U2OS, MEF, CHO-K1, NHEK and NHDF cells, which were immortalized from donors. In all of the cells, we observed the same principal frequency of 0.004 min^-1^ with the higher harmonics (Fig. 1C). The average amplitude of the FT components and the 1/f noise varied depending on the cell type, but the frequency spectrum remained strikingly similar.

The estimated amplitude of the dry mass change is approximately 100 pg (CHO-K1 cells) that exceeds three times method precision (Fig. S4, S5). We made several tests to rule out the possibility that the oscillations might have been experimental artifacts. First, we looked for and found no periodicity in the experimental conditions, such as the incubator temperature (Fig. S6A). We next varied the number of cells and performed the same FT analysis. The characteristic frequencies, which are distinct from the 1/f noise, could be found only when more than 100 cells were analyzed (Fig. S6B). This confirms that the rhythm depends on the cells. In particular, the cells must be alive, as no oscillations were found when the cells were fixed (Fig. S6B, lower right). Finally, the FT analysis of a random dataset failed to detect any periodicity, confirming that the rhythm was not a result of the mathematical analysis (Fig. S6C).

### Role of Cell Cycle and Circadian Oscillators

We have revealed a periodicity (4 hr rhythm) in cell dry mass dynamics that seems to be general in mammalian cells. A priori, this could be an ultradian rhythm with a period of 4 hours, a harmonic of a longer-period biological clock, or an oscillator such as circadian rhythm or cell cycle.

As a next step, we investigated potential links between the 4 hr rhythm and the cell cycle. Cell cycle synchronization did not significantly change the period or the amplitude of the rhythm (Fig. S7A). Despite the variability in cell cycle time, ranging in our experiments between 12.1h and 21.3h, all analyzed cell lines/types showed the same oscillation period of 4 hours (Fig. 1E), suggesting that the rhythm was not a cell-cycle harmonic. Furthermore, varying the temperature from 33°C to 39°C significantly influenced the cell cycle length in HeLa and CHO-K1 cells, while leaving the period of dry mass oscillation unchanged (Fig. 2B and Fig. S7B). Thus, in contrast to the cell cycle, the 4 hr rhythm showed perfect temperature compensation, a hallmark of a biological clock *(3, 4)*.

**Fig. 2.**
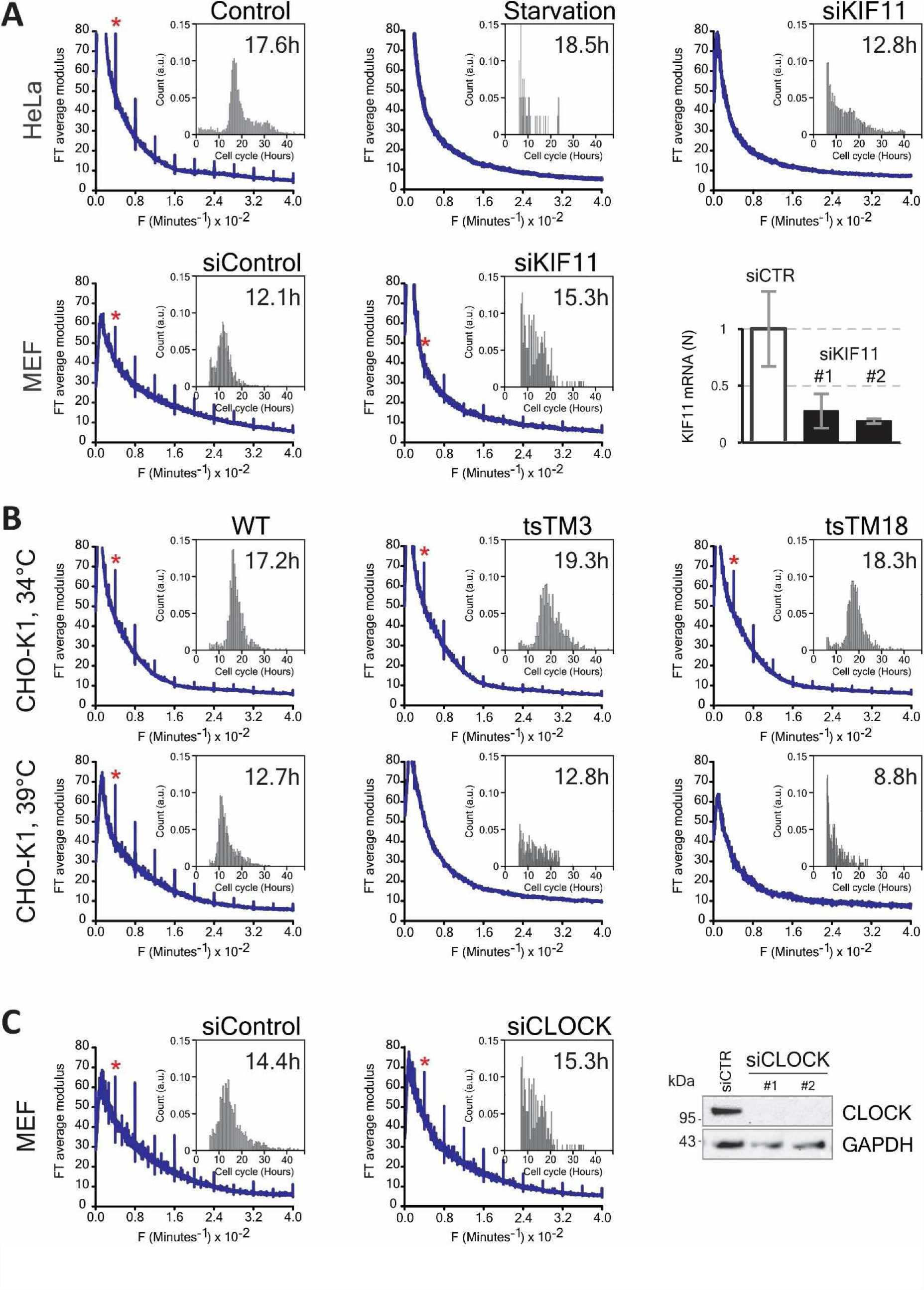
(A) Effect of cell cycle inhibition on the 4 hr rhythm: serum starvation and siKIF11 treatment were performed in HeLa (upper row) and MEF (lower row) cells. Quantitative PCR analysis (right hand plot of the lower row) shows the inhibition of KIF11 expression by two specific siRNAs in MEF cells. The inserts show the distribution of the corresponding cell cycle times. (B) Effects of cell cycle inhibition are shown in temperature-sensitive CHO-K1 mutants at 34°C (upper row) and 39°C (lower row). In the insert, the distribution of the cell cycle times is shown, and their mean values are in hours. (C) Effect of circadian rhythm inhibition by siCLOCK is shown in MEF cells. The blots show the inhibition of CLOCK expression by siCLOCK1 and 2 in MEF cells. Inserts represent the distribution of the cell cycle times.

To further examine the role of the cell cycle, we used serum starvation to induce G0/G1 cell cycle arrest in HeLa cells. Over 48 hours of starvation, the cells gradually ceased proliferating (Fig. S7C). Curiously, even though the cells were still alive, the 4 hr rhythm disappeared (Fig. 2A and Fig. S7C). When HeLa cells were arrested in the G2/M phase by knocking down the KIF11/EG5 gene, which encodes a motor protein required for mitotic spindle formation, the rhythm was also suppressed (Fig. 2A and Fig. S8A, movie S1). By contrast, in MEF cells, where the mitotic arrest induced by KIF11 knockdown was not as efficient, the rhythm was still observable (Fig. 2A, movie S2).

Next, we examined the temperature-sensitive CHO-K1 cell lines tsTM3 and tsTM18, which carry mutations in ubiquitin-activating enzyme UBA1 and DNA replication regulator SMU1, respectively (Fig. 2B). These mutants have been shown to undergo S-G2 and S and G2 cell cycle arrest, respectively, at the nonpermissive temperature of 39°C *(5-8)*. We found that at the permissive temperature of 34°C, the tsTM3 mutant displayed a 4 hr rhythm comparable with that of control CHO-K1 cells. In tsTM18 cells, oscillations were also detected, albeit with a reduced amplitude. At 39°C, however, the rhythm was suppressed in both mutants, but it was still observed in wild-type CHO-K1 cells (Fig. 2B).

These results show that cell cycle arrest at any stage suppresses the 4 hr rhythm. Therefore, even though the dry mass oscillations we observed were not a direct consequence of the cell cycle, the oscillations seem to be a feature of proliferating cells.

Because 4 hr is an integer submultiple of 24 hr, and, similar to the circadian rhythm, the 4hr-rhythm is temperature-compensated; therefore, the 4 hr rhythm might have been a circadian harmonic. However, all transformed cell types we examined showed a robust 4 hr rhythm (Fig. 1C), whereas circadian rhythms have been documented only in U2OS and MEF cell lines *(9)*. Finally, knocking down CLOCK, a major/master transcriptional regulator of circadian rhythms, in MEF and U2OS cells did not affect the dry mass oscillations (Fig. 2C and Fig. S8B for U2OS).

Altogether, these findings suggest that the periodic changes in cell dry mass represent a new ultradian rhythm in growing cells.

### Role of Cell Machineries

Proteins represent more than half of the cell dry mass, and that is followed by nucleic acids (∼15%), lipids (∼10%) and sugars *(10, 11)*. All of these constituents have approximately the same specific refractive index increment (∼0.18 µ m^3^/pg) and contribute to the measured dry mass in proportion to their abundance. Therefore, the periodic change in the dry mass we observed might result from the synthesis/degradation of these molecules or from their transport into or out of the cell.

To obtain more information on the molecular mechanisms controlling dry mass oscillations, we used: (i) actinomycin D (ActD) to inhibit transcription, (ii) cycloheximide (CHX) to inhibit protein synthesis, and (iii) proteasome inhibitors MG132 and bortezomib (Btz) to block ubiquitin-dependent protein degradation. We verified that the drugs were used at concentrations that did not induce excessive cell death over a 48-hr observation period (Fig. S9). To analyze the drug response, we used the average Fourier amplitude of the fundamental peak (AF) to define the oscillator strength. This parameter showed very little variability within a cell batch, providing a reliable and quantitative measure of the inhibition effect (Fig. S10). Since we have shown that the 4 hr rhythm is a feature of proliferating cells (Fig. 2A and B), its inhibition by the drugs could be either direct, or could be caused by cell cycle arrest. To exclude the latter effect, we analyzed only the cell trajectories between two well defined mitotic events, thus ensuring that the cells treated by the drug were still cycling (or able to undergo at least one mitosis).

We observed that all the drugs decreased the strength of oscillation to varying degrees in MEF and HeLa cells (Fig. 3A). Significantly, the periodicity, which is the defining feature of the ultradian clock *(3, 4)*, remained unchanged. Transcription inhibition with ActD failed to arrest the 4 hr rhythm in MEF cells, even at the highest dose that induced ∼50% cell mortality (Fig. 3A and Fig. S9). In contrast, inhibition of either protein synthesis (CHX) or degradation (MG132 and Btz) completely suppressed the rhythm and exhibited relatively low cytotoxicity (Fig. 3A and Fig. S9). Similar effects were observed in HeLa cells where low doses of CHX or Btz (few % cell mortality) completely eliminated the 4 hr rhythm, while ActD did not show any significant AF decrease at low doses (Fig. 3A). Note that the doses of MG132 required to arrest the 4 hr rhythm in HeLa cells were also significantly cytotoxic (∼50% cell death, Fig. S9).

**Fig. 3.**
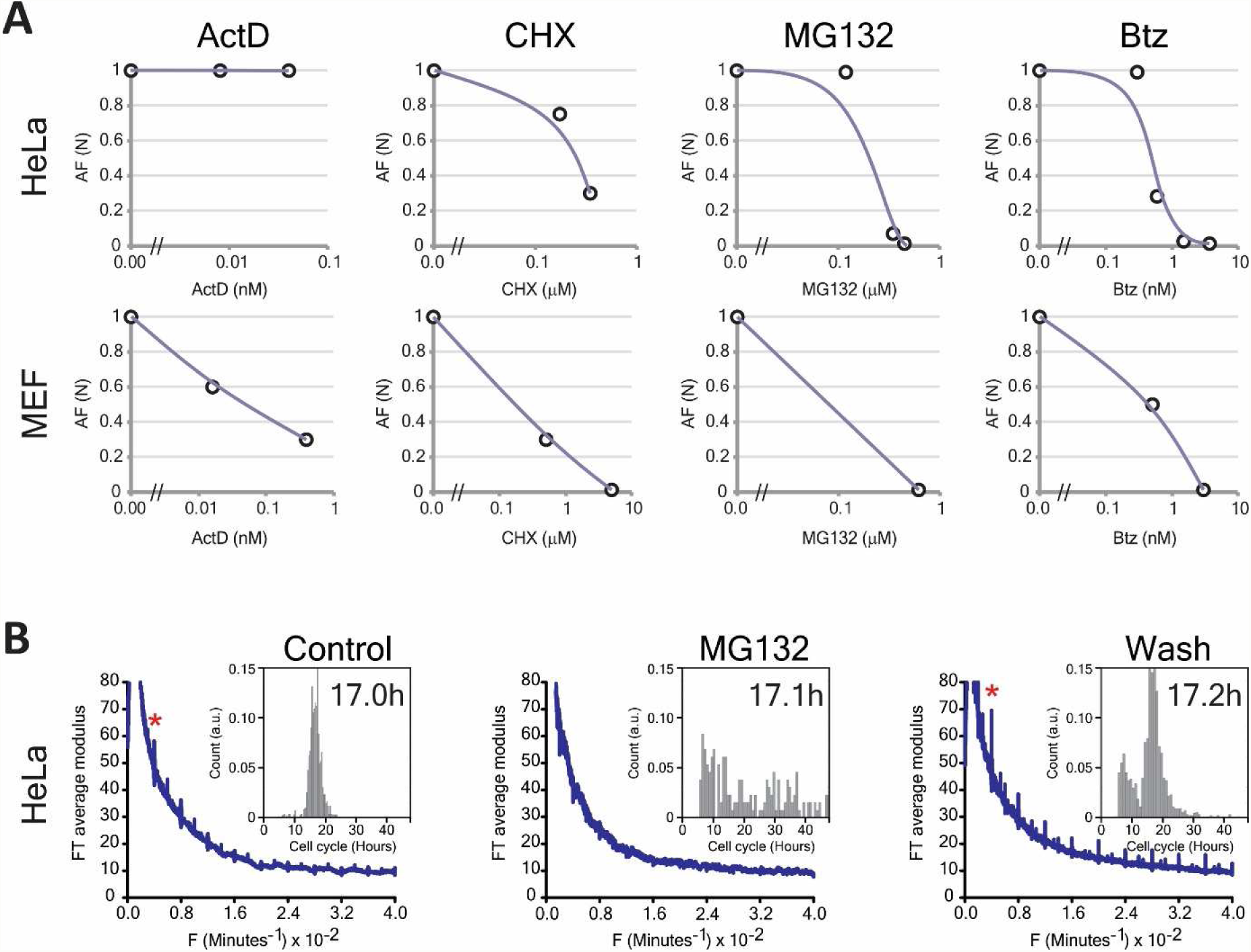
(A) Effect of different inhibitors on the rhythm were measured by the amplitude of the Fourier transform of the fundamental peaks after subtraction of the 1/f noise. The data are normalized to the control values. (B) Inhibition of the 4 hr rhythm by MG132 is reversible (see text for full explanation). Inserts represent the distribution of the cell cycle times.

These results show that the oscillations of the cell dry mass are particularly sensitive to proteostasis inhibition and therefore may represent the periodic changes in protein level during cell growth. The effect of the proteasome inhibitor Btz was the most notable. In HeLa cells, significant suppression of the 4 hr rhythm (without visible cytotoxicity) occurred with 0.6 nM Btz treatment, which is the Ki value for its specific target: the 20S proteasome (Fig. 3A) *(12)*. Using an alternative proteasome inhibitor MG132 at higher doses of 0.45µ M also suppressed the rhythm, and we established reversibility in its action, by showing that subsequent removal of MG132 by rinsing restored the oscillations. Strikingly, the restored 4 hr rhythm had a greater strength than it had before the treatment, suggesting that the transient proteasome inhibition either increased the amplitude of protein pulses or induced more cells to oscillate (Fig. 3B). Therefore, our results demonstrate that the proteasome is one of the main molecular machineries regulating the 4 hr ultradian rhythm.

## Conclusions/Discussion

In conclusion, we document a new ultradian rhythm that occurs in a wide range of mammalian cells during the interphase of the cell cycle. The rhythm has a period of 4 hours and was observed by holographic lens-free microscopy by measuring the individual histories of the dry mass of thousands of live cells, with a resolution of five minutes. We emphasize that these results were obtained using inherently heterogeneous cell populations, thus eliminating synchronization and labeling bias. The 4 hr rhythm is suppressed by proteostasis disruptors and is detected only in proliferating cells, suggesting that it represents the periodic dynamic of the protein mass in the growing cell.

Periodic changes in protein levels have been shown previously; the best known example is the oscillations of the key regulators of the cell cycle *(10, 13, 14)*. Recent proteomics studies suggested that the abundance of more than 10% of cellular proteins are subject to the 24 hr circadian rhythm *(15)*. Higher frequency ultradian oscillations have also been reported. In the early 1960s, Brodsky and collaborators, by analyzing the incorporation of radioactively labeled amino acids into cellular proteins, suggested a 1-2 hr periodicity in the protein synthesis rate *(16)*. The same authors have reported the observation of ultradian dry mass oscillation in retinal ganglion cells, as observed by interferometry *(16)*. In the last decade, ultradian oscillators originating in transcriptional-translational-posttranslational feedback loops have been discovered for transcription factors p53, Hes1, and NFkB *(17-19).* Notably, there was a change in the level of the proteins implicated in these signaling circuits, which have periods between 2 and 6 hours. More recently, the oscillations of another transcriptional regulator, XBP1, has been shown to coordinate a new 12 hr ultradian rhythm *(20)*. Very recently, Liu et al. *(21)* reported repetitive dips in the coefficient of variation (CV) of the cell growth rate in HeLa cells. The authors used quantitative phase microscopy interferometry to measure the dry mass of the cells during the cell cycle at a 30 min time resolution. They tentatively suggested that the dips in the cell growth rate CV might reflect a novel oscillatory circuit in protein synthesis/degradation that is intrinsic to cell growth rate regulation. Although the reported periodicity was close to 4 hours, it was significantly temperature dependent: 4.7 hr at 33°C and 5.8 hr at 36°C.

The 4 hr rhythm we describe here differs, however, in several important ways from the previous observations. First, it appears to be cell-autonomous, robust and universal, as it was found in all cultured mammalian cells we examined. The rhythm is present in asynchronous cell cultures growing in standard conditions, and no additional stimuli are required to trigger it. Second, the 4 hr rhythm is not limited to a few specific proteins; rather, it involves global changes in the total mass of cell constituents. Third, the 4 hr rhythm is temperature–compensated; this was not the case for the ultradian rhythms mentioned above. Finally, our analysis with the inverse Fourier transform indicates that the 4 hr rhythm has a particular nonsinusoidal waveform, where the long delay periods are followed by rapid (∼30 min) symmetric changes in the cell dry mass (Fig. 1B and Fig. S5). This pulsatile dynamics may explain why the rhythm was not observed in previous works that used lower time resolution (>30 min) in sampling. We also needed to follow hundreds of cells in parallel to begin to see this periodic signal, which required quantitative phase imaging techniques with a large field of vision.

Our results give a first glimpse into the underlying mechanism of the 4 hr oscillator. The rhythm disruption by proteasome inhibition and its stimulation upon inhibitor removal suggests that the proteasome is implicated in oscillator regulation. This is not surprising, as the proteasome degrades key pacemaker proteins, meaning that it has an essential role in almost all reported biological rhythms. The universality and the amplitude of the mass oscillations we see (Fig. 1 and Fig. S5) suggest that the proteasome, by itself, is a 4 hr rhythm pacemaker, and its activity is responsible for pulsatile dynamics of the total mass of proteins. The existence of posttranslational proteasome-based oscillators has been predicted previously by mathematical models that comprise both protein synthesis and degradation *(22-25)*. It should be noted that our analysis cannot determine whether the dry mass is rising during the pulses as a result of increased synthesis or is dropping because of accelerated degradation (Fig. 1B). Even though both possibilities remain, the second hypothesis seems more thermodynamically likely.

Another aspect of the 4 hr rhythm is that it is linked to the cell cycle. Curiously, pioneering work by Klevecz suggested that endogenous oscillations in protein synthesis set the generation time of the cell cycle as a multiple of a fundamental 4 hr period *(26, 27)*. This ultradian oscillator was found to be temperature compensated *(26)*. Lloyd and Volkov *(28)* later proposed a mathematical model for the cell cycle to explain these results. This model included a fast ultradian component with pulsatile dynamics that were similar to what we see in our reconstructed signal *(28)*. Therefore, the 4 hr rhythm we describe here may reflect the same timekeeping mechanism that regulates cell growth during interphase and dictates cell cycle duration. Although we found that the inhibition of the 4 hr rhythm with low doses of proteasome inhibitors did not block ongoing cell division, it certainly prevented subsequent mitosis because the cultures finally stopped growing (Fig. S11). Conversely, the suppression of the 4 hr rhythm following cell cycle arrest suggests a strong coupling of the two oscillators. Finally, as the proteasome is the ultimate regulator of cyclin/CDK machinery, our conclusions seem to be potentially compatible with the current models of cell cycle regulation.

## Supporting information

MaterialandsupplFigures_lGhenim

## Acknowledgments

L.G. is very grateful to Dr. Kimihiko Sugaya of the Research Center for Radiation Protection, National Institute of Radiological Sciences, Chiba, Japan for making the TSTM3, TSTM18 and CHO-K1 cells available to us. We also thank Dr. D. A. Skoufias from IBS (Institut de Biologie Structurale, (VIC)-Cell Division Team), Grenoble for providing HeLa cells. The company IPRASENSE (Clapiers, France) and Dr. F. Navarro (Department of micro and nano Technologies for Healthcare and Biology, LETI CEA Grenoble) generously lent us the lens-free microscopes. C.A is very grateful to Yves Usson (TIMC-IMAG, Uni. Grenoble Alpes, CNRS UMR 5525), Julien Savatier and Serge Monneret (Aix Marseille Univ, CNRS, Centrale Marseille, Institut Fresnel, Marseille, France) who contributed to the quantitative phase imaging comparisons. C.A is very thankful to Romaric Vincent and Thomas Bordy, for their help in data acquisition and data analysis.

## Author contributions

Original Concept: X.G; Experimental Design: L.G; M. B. and X. G.; Experiments carried out by L.G. with help of P.O. (PCR, Western blot, dose responses); Image Reconstruction, total phase calculation and cell cycle identification by L. H. and C.A.; Cell tracking by L. G.; Data analysis: Fourier and statistical analysis: L. G. and J. Y. F.; The manuscript was prepared by L. G.; M. B. and X. G.

## Data and materials availability

Data are available in the main text and in the supplementary materials. Image and trajectory data can be supplied by request to the authors.

## Supplementary materials

Material and Methods

Figures S1-S11, movies S1, S2

