## Supplementary material for "A New Ultradian Rhythm Linked to Protein Degradation and Synthesis in Mammalian Cells": MaterialandsupplFigures_lGhenim

**Materials and methods**

**Cell culture**

*HeLa cells* were cultured in high glucose DMEM supplemented with GlutaMAX, pyruvate, and 10% calf serum (Gibco). *NHEK cells* (Lonza) were cultured in KBM-gold Bulletkit (Lonza) and were passaged with the ReagentPackSubculture reagents (Lonza). NHDF-Ad-Der Fibroblasts FGM-2 cells (Lonza) were cultured in FGM-2 BulletKit (Lonza) and were passaged with the ReagentPack Subculture Reagents (Lonza). *U2OS cells* (ATCC) were cultured in McCoy’s 5A medium (ATCC) and 10% calf serum. Mouse Embryonic Fibroblasts (C3H/10T1/2 clone 8, ATCC) were cultured in Eagle’s Minimum Essential Medium (EMEM; ATCC) and supplemented with 10% calf serum. *CHOK1-WT, TSTM3* and TSTM18 (Research Center for Radiation Protection, Chiba, Japan) were cultured in Nutrient Mixture F-12 Ham (SIGMA) supplemented with 1mM L-glutamine (SIGMA) and 10% calf serum. All media was supplemented with 1% Penicillin /Streptomycin (Invitrogen).

**Cytotoxicity assays: determination of LD50 by dose response:**

LD50 was determined by testing in duplicate the drugs at different concentrations on cells seeded 24 hours, in 96-well plates, prior to drug treatment. CellEvent™ Caspase-3/7 Green Detection Reagent (Invitrogen, Ref. C10423, C10723) was added, to each well, directly with the drug mixture in a final concentration of 2 µM per well. 24 and 48 hours after drug and treatment with the CellEvent™ Caspase-3/7 reagent, cells were counterstained with Hoechst. Images were acquired using a High Content CellInsight™ NXT HCA automated microscopes (Thermo Scientific™). Analysis was performed using R software and the percentage of dead cell was evaluated based on negative control levels. The real time imaging experiments with cells treated with different drugs were then performed during 48 hours at concentrations much below the concentration that induces 50% mortality.

**Drug washout experiments**

All drugs were prepared first in DMSO (Sigma-Aldrich). The cells were treated with drugs 24 hours after being seeded and grown at 37°C. To inhibit transcription, Mouse Embryonic Fibroblasts and HeLa cells were treated respectively with 0.016 nM, 0.4 nM and 0.008nM, 0.04 nM of Actinomycin respectively. To inhibit translation HeLa cells and Mouse Embryonic Fibroblasts were treated respectively with 0.175 µM, 0.35 µM and 0.5 µM, 5µM of Cycloheximide respectively. To inhibit protein degradation HeLa and MEF were treated respectively with 0.3 nM, 0.6 nM, 3.75 nM, 5 nM and 0.5 nM, 3nM of Bortezomib. Otherwise, to inhibit protein degradation HeLa and MEF cells were treated respectively with 0.12 µM, 0.35 µM 0.45 µM, and 0.62 µM of MG132. To reverse the inhibition of protein degradation, HeLa cells were washed out three times with a medium without drugs.

**Small interfering RNAs Transfection**

RNAi screens were performed in 6-well plates and three Fibronectin (F1141-5MG from Sigma_Aldrich) coated 35 mm glass bottom dishes (ref 150682 Thermo Scientific). RNA interference against EG5 (ref Qiagen Hs_KIF11_6 Cat N°./ID: SI02653693, Mm_Kif11_3 FlexiTube siRNA (5 nmol) Cat No./ID: SI01082389, Mm_Kif11_2 FlexiTube siRNA Cat No./ID: SI01082382), and against Clock (Mm_Clock_6 FlexiTube siRNA Cat N°./ID: SI02707908; Mm_Clock_2 FlexiTube siRNA Cat N°./ID: SI00168595) were performed with two independent siRNAs from Qiagen each, and each siRNA transfection was repeated two times. As control we used siRNA AllStar negative control (AS) (Qiagen, Ref. 1027281). HeLa and C3H10T1/2 cells (20 000 cells per petri dish and 80 000 cells per well) were seeded or plated in complete medium 24 hours before transfection. Transfection of these cells was performed using Lipofectamine® RNAiMAX reagent. The RNA-lipid complexes were prepared and then added to the cells. First, Lipofectamine® RNAiMAX Reagent was diluted in Opti-MEM® Medium, second siRNA was diluted in the same solution. The two mixtures were incubated at room temperature for 10 minutes. Diluted siRNA was added to the diluted Lipofectamine® RNAiMAX Reagent and this mixture was then well homogenized and incubated for 15 minutes at room temperature. Finally, the siRNA-lipid complex was distributed on the cells. The final concentration of siRNA per well was 1 to 10 nM. The plates as well as the glass bottom dishes were placed in the incubator at 37°C for 48 hours. The glass bottom dishes were placed on the lens-less microscopes for data acquisition within the incubator. The transfection was stopped after 48 hours. The 6-well plates were used for the PCR and western blot experiments as controls of the optical experiments.

**Western Blot analysis**

After 48 hours in the incubator, cells in the 6-well plates were lysed using RIPA buffer (Sigma, Ref. R0278) supplemented with Complete^TM^ Mini Protease Inhibitor Cocktail (Sigma, Ref.P8340). The buffer (50 µl) was added to each well and the plate was incubated at 4°C for 30 minutes. The lysates are homogenized, collected and put in a 1.5 mL Eppendorf tube at 4°C for another 15 minutes. Centrifugation was then done at 13000 rpm for 15 minutes at 4°C and the supernatant was taken and stored at -20°C while waiting for the protein dosage. The protein dosage was done according to Pierce™ BCA Protein Assay Kit protocol (Thermo Scientific, Ref.23227). The range of BCA was made from a commercial solution at 2 mg/ml. Dilutions are made in RIPA. Quantification was done in duplicates with 5 µl samples and 100 µl BCA mix prepared according to the manufacturer’s protocol. The plate was then incubated for 45 minutes at 37°C and the reading of the plate was performed using the TECAN Infinite® M1000 reader at 562 nm wavelengths. To examine the expression of the target genes, the same amount of total protein was analyzed by western blot using target-specific antibodies. For siEG5 and siClOCK experiments, 7µg (protein samples) were subjected to polyacrylamide gel electrophoresis on NuPAGE™ Novex™ 12% Bis-Tris Protein Gel (Life Technologies, Ref.NP0342BOX) in MES buffer. The separated proteins were transferred onto a nitrocellulose membrane (D. Dutscher, Brumath, France) using iBlot™2 Transfer Stack according to the manufacturer’s protocol. The membranes were blocked in 5% non-fat milk in Tris-buffered saline, 0.1% Tween 20 (TBST, Sigma) for 1 h at room temperature (RT) and incubated with primary antibodies diluted at 1:500 in 5% non-fat milk in TBS-T overnight at 4°C. We used rabbit polyclonal antibodies directed against *human EG5 (1/1000,* ab171963), rabbit polyclonal antibodies directed against *human and mouse CLOCK (*1/2000, ab93804), mouse polyclonal antibodies against tubuline (1/2500, SIGMA T-6074) and actin (1/5000, (H-196)) Santa Cruz Biotechnology Sc-7210 and rabbit polyclonal Anti-GAPDH ((FL-335) Santa Cruz Biotechnology Sc-25778). Membranes were washed three times for 10 min in TBS-T and incubated with secondary horseradish peroxidase-conjugated antibodies (anti-mouse or anti-rabbit, eBiosciences, Paris, France) at a dilution of 1:10 000 in TBST for 1 hour at room temperature. Membranes were washed three times for 10 min in TBST. Detection was performed with an ECL Prime kit [SuperSignal™ West Pico PLUS Chemiluminescent Substrate (Thermo Scientific, Ref.34577) and the SuperSignal™ West Femto Maximum Sensitivity Substrate (Thermo Scientific, Ref.34095) according to the manufacturer’s guidelines.,]. Chemiluminescence was analyzed using a ChemiDoc Touch Imaging System (BioRad, Marne-la-Coquette, France), and quantification of the intensity of the bands was performed using the ImageLab™ Touch Software (BioRad).

**Gene expression analysis by RT-qPCR**

The commercial RNeasy® Plus Mini Kit (Qiagen, Ref.74136) was used for total RNA extraction according to the manufacturer’s instructions. 48 hours after the Embryonic Mouse Fibroblasts (C3H/10T1/2 clone 8) (as above for invasion experiment) transfection with the siRNA EG5 (KIF11-2 and KIF11-3), the media was removed and the cells were lysed directly in the wells with 350 µl Buffer RLT supplied by the kit. The total RNA was eluted in 40 µl of RNase-free water and the sample were stored at -80 °C.

RNA concentration was measured using a Thermo Scientific NanoDrop™ 1000 Spectrophotometer (Thermo Scientific). Then, 2 µg RNA were reverse-transcribed transcribed in a total volume of 20 μl using SuperScript™ IV VILO™ Master Mix (Invitrogen, Ref.11756050) according to the manufacturer’s instructions. The cDNA was stored at −80°C until further use. Reverse transcription reactions were diluted to 1/10 in RNAse free water and 3µl of the diluted cDNA was used for each quantitative PCR. qPCR was carried out using sequence-specific primers on a StepOnePlus™ RealTime PCR System (Applied Biosystems /Thermo Fisher Scientific) in 96-well plate with SYBR® Green dye (Platinum™ SYBR™ Green qPCR SuperMix-UDG, Thermo Scientific, Ref. 11733-046). All experiments were run in triplicate, and the results were normalized to GAPDH gene (mKIF11-F1 and R1, Ref 804490-689.15 and mGAPD-HF1 and R1, Ref 804490-955.34, Eurogentec).

**Lens-free microscopy and holographic reconstruction**

Lens-free acquisitions were performed with a Cytonote microscope (Iprasense, Montpellier, France). This setup is inspired by the lens-free imaging system described in *(1)*, which was modified to perform continuous monitoring of cellular cultures inside an incubator at a controlled temperature and humidity *(2)*. Illumination is provided by a red LED (647 nm, FWHM 13 nm). The light passes through a 150 µm pinhole at a distance of approximately 5 cm from the sample. The CMOS sensor is in contact with the cell culture recipient at a distance of ∼1.5 mm (see Fig. S1).

In the absence of optics, the sample image is obtained by a reconstruction algorithm. For lens-free microscopy, the reconstruction problem is usually formulated under the first Born approximation *(3)*, which restricts the solutions to low scattering objects. Under this approximation light transport is described by the Fresnel propagator which can be analytically inverted. This propagator applied to the field of light in the sensor plane allows estimating the field of light just after the object plane. However the reconstruction problem remains ill-posed since the lens-free setup, in the absence of a reference arm, does not record completely the field of light in the sensor plane but only its intensity. Yet the lack of phase information does not prevent from retrieving the sample image. The problem can be tackled with the following inverse problem approach. Light is described as a complex scalar field and its propagation from one plane to another is obtained using a convolution kernel *h_Z_*. Therefore, if *A_0_* is the complex field after the sample, the field *A_Z_* in the detector plane at the distance *Z* from the object is:

$A_{0}=A_{Z}*h_{Z}$	(1)

where ‘*’ is the convolution operator. Here, we used a kernel *h_Z_* derived from Fresnel theory:

$h_{Z}=\left( \frac{1}{iZ} \right)e^{\frac{i\vec{r}^{2}}{\lambda Z}}$	(2)

i being the imaginary unit and $\vec{r}$ the spatial position in the horizontal plane. In the following, Z is assumed to be known. In practice, it is estimated with a precision of approximately 10 μm by inspecting Z-stack of back-propagated measurements. The convolution of Eq. (1) can be explicitly written as:

$A_{Z}\left( \vec{r}^{'} \right)=\int d\vec{r}A_{0}\left( \vec{r} \right).h_{Z}\left( \vec{r}^{'}-\vec{r} \right)$	(3)

The quantity measured by a standard detector is the intensity of the light field: $I_{Z}=\left| A_{Z} \right|^{2}$, where the modulus of $A_{z}$ is known but not its phase $\varphi_{Z}$. Whatever the phase $\varphi_{Z}$, $A_{Z}=\sqrt{I_{Z}}.e^{i\varphi_{Z}}$ and the corresponding field $A_{0}$ at the sample plane, $A_{0}=\left( \sqrt{I_{Z}}.e^{i\varphi_{Z}} \right)*h_{-Z}$ would perfectly match the data. The inverse problem approach consists here in finding the phase $\varphi_{Z}$ so that $A_{0}\left( \varphi_{Z} \right)$ minimize the total variation (TV) criterion $\epsilon\left( \varphi_{Z} \right)$:

$\epsilon\left( \varphi_{Z} \right)=\int d\vec{r}\sqrt{\varepsilon+\frac{\delta A_{0}\left( \vec{r} \right)}{\delta x}+\frac{\delta A_{0}\left( \vec{r} \right)}{\delta y}}$	(4)

Where ε is a small valued coefficient (10^-4^) used to make Eq. (4) differentiable and prevent division by 0 in the gradient derivation presented afterwards. The second term is a *L_1_*-norm applied to the fields $\frac{\delta A_{0}\left( \vec{r} \right)}{\delta x}+\frac{\delta A_{0}\left( \vec{r} \right)}{\delta y}$ which promotes sparsity. The optimization is performed using the non-linear conjugate gradients iterative method as described by the following pseudocode:

1/ initialization $k\leftarrow0, D^{(k)}\leftarrow0, \varphi_{Z}^{\left( k \right)}\leftarrow0$

2/ $k\leftarrow k+1$

3/ computation of gradient $\nabla\epsilon^{\left( k \right)}$, of advancement direction $D^{(k)}$and advancement step $\sigma^{(k)}$

4/ phase update: $\varphi_{Z}^{\left( k \right)}\leftarrow\varphi_{Z}^{\left( k-1 \right)}+\sigma^{(k)}.D^{(k)}$

5/ test of convergence, if not reached go back to 2/

The advancement direction $D^{(k)}$ is obtained by conjugating the gradient $\nabla\epsilon^{\left( k \right)}$ with the previous direction of advancement$D^{(k-1)}$, using Hestenes-Stiefel formula (4). The advancement step $\sigma^{(k)}$ is obtained by developing the TV criterion at order 2 in the advancement direction (Hessian computation) and by minimizing a second-order polynomial of variable $\sigma^{(k)}$. For the test of step 5/, we simply stop the algorithm when the number of iterations reaches 15.

Fig. S2 illustrates the holographic reconstruction process described here above. Fig. S2a shows the full field of view of a raw acquisition of culture of HeLa cells. Based on this image the holographic reconstruction produces a phase image of the cells as shown in Fig. S2c. The reconstructed phase image exhibits locally wrapped phase artefacts corresponding to cells that induce a phase shift exceeding +π. In order to perform a simple phase unwrapping, we detect the contiguous pixels with phase value below 0, and set these pixels to +π (Fig. S2e).

**Cell-tracking**

We used the Trackmate algorithm, an open source Fiji plugin for the automated tracking of single particles *(5).* The Trackmate Fiji plugin guides the user through several stages of the cell-tracking algorithm, namely a cell detection stage, a cell tracker stage and several filters applied to the cell detections and the computed tracks. We used the following parameters: the estimated blob diameter was set to ~15 pixels, the detector threshold was set to 0.7, the linking maximum distance was set to 15 pixels (~25 µm), the gap-closing max distance was set to 15 pixels, gap closing max frame gap to 5 and the filter number of spots on tracks was set to 10. At the end of the cell-tracking process, results are generated in the form of text files. The output file ‘Spots in tracks statistics’ contains a table listing all detected cells with their respective positions in the acquisition, their frame number and their track number.

**Cell dry mass calculation**

The phase recovered from lens-free microscopy is proportional to the density and thickness of the specimen layer *(6)*. A relation is defined between the phase shift and the optical path difference (OPD), which quantifies the integral of the sample object refractive index difference with respect to the surrounding medium along the optical path:

$\varphi_{shift}\left( x,y \right)=\varphi\left( x,y \right)-\varphi_{medium}$	(5)

$$OPD\left( x,y \right)=\lambda.\frac{\varphi_{shift}\left( x,y \right)}{2\pi}\int_{0}^{h} \left[ n\left( x,y,z \right)-n_{medium} \right]\delta z$$

where $n$ is the local sample refractive index, $n_{medium}$ is the is the surrounding medium refractive index, z is the position along the optical axis, h is the thickness of the sample object and λ is the illumination wavelength. The *OPD* values can be integrated over the total projected area *S* of the cell. Here we used the optical volume difference ($OVD$) denomination introduced in *(7)* to define this integral. It is expressed as a unit of volume in μm^3^:

$OVD=\int_{S} OPD(x,y)\partial x\partial y$	(6)

To determine the cell area *S*, we used a seeded growing segmentation algorithm controlled by a single parameter, a phase threshold value which delineates the separation between the background and the cell area (see Fig. S3). A relationship between the phase shift measurement and the cell mass has been defined in *(8, 9)* and can be used to convert $OVD$ to cell dry mass ($CDM$) measurements, the mass of the cellular content except water. Under our notation, this relationship is simply given by:

$CDM=\frac{OVD}{\alpha}$	(7)

Where α is the specific refractive increment which relates the refractive index change to increase in mass density. There is a variety of substances within a cell. However the specific refractive index of these substances falls with a narrow range and Barer *(8)* defined an α constant of 0.18 µm^3^.pg^-1^ for most eukaryotic cells, taking into account not only proteins, but also lipids, sugars, and nucleic acids

To estimate the precision of the *CDM* measurement obtained by means the lens-free microscope we compared the cell measurements with two other techniques: digital holographic microscopy (DHM) and quadriwave lateral sheering interferometry (LSI). The measurements obtained with lens-free microcopy correlate linearly with the measurements obtained with DHM and LSI (see Fig. S4). It suggests that the lens-free microcopy setup can be considered as a quantitative phase imaging technique for the measurements of adherent cells. The values measured with lens-free microcopy are however under-estimated, they must be corrected by a factor of ~1/0.65. This is due to the sparsity constraints used in the holographic reconstruction algorithm (See Eq. 4) which reduces overall the phase signal. According to a methodology and the dataset described in *(6)*, we can estimate the precisions of the *CDM* measurements obtained with the lens-free microscopy setup used in this study to be about 35 pg.

**Fourier Transform and other spectral analysis algorithmes**

The Fourier transform algorithm allows real-time spectral analysis of the dry mass and help us to find the spectrum of possible periodic signals. We compute the Fourier transform over the duration T(j) of our measurement of the dry mass of each cell j during interphase. T(j) is defined to be the time between the two local spikes ***(2)*** of dry mass that correspond to the successive divisions that delimit the interphase or, for incomplete cycles, the time from the first local spike of dry mass until tracking of the cell is lost. After removing the background, the signal was convoluted with a sine window function that vanishes at the limiting times. The FT was then performed for each frequency with an increment equal to df(j) = 1/(T(j)*M) depending on cell j, where M is an integer corresponding to the chosen resolution in frequency sampling. We fix M to be equal to 25 for all the samples. The set of all transforms was analyzed statistically by computing the averaged absolute amplitude, as well as the standard deviation, in each frequency bin df= n_f_ /(T_max_*M). The averaged FT was performed using bins defined by the longest time duration T _max_ = max{T(j)}, and n_f_ is an arbitrary positive integer, defining the size of the bin. In general, we take n_f_ =1 or 2. The averaged absolute amplitude was properly normalized in order to take into account the different bin sizes and lengths of the traces.

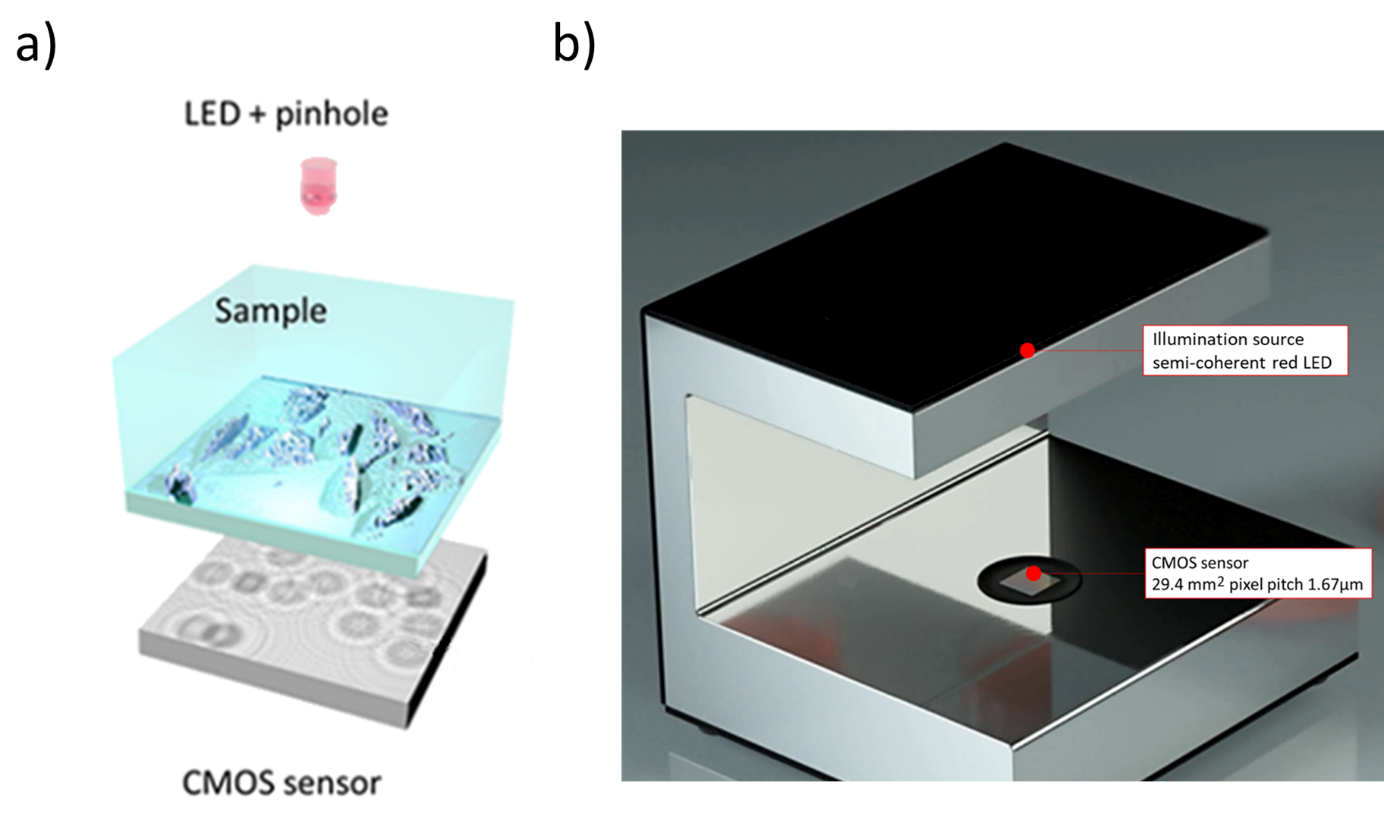

**Supplementary Fig. S1. Lens-free microscopy** (a) Schematic of the lens-free microscopy acquisition setup. The CMOS sensor is put in contact with the Petri dish and records the holographic pattern resulting from the interference between the partially coherent incident light and the light scattered by the cells. (b) The lens-free microscope is the Iprasense Cytonote. It features a CMOS image sensor with a pixel pitch of 1.67 μm and an imaging area of 6.4 mm x 4.6 mm. Illumination is provided by a red LED along with a 150 µm pinhole placed at a distance of approximately 5 cm from the sample.

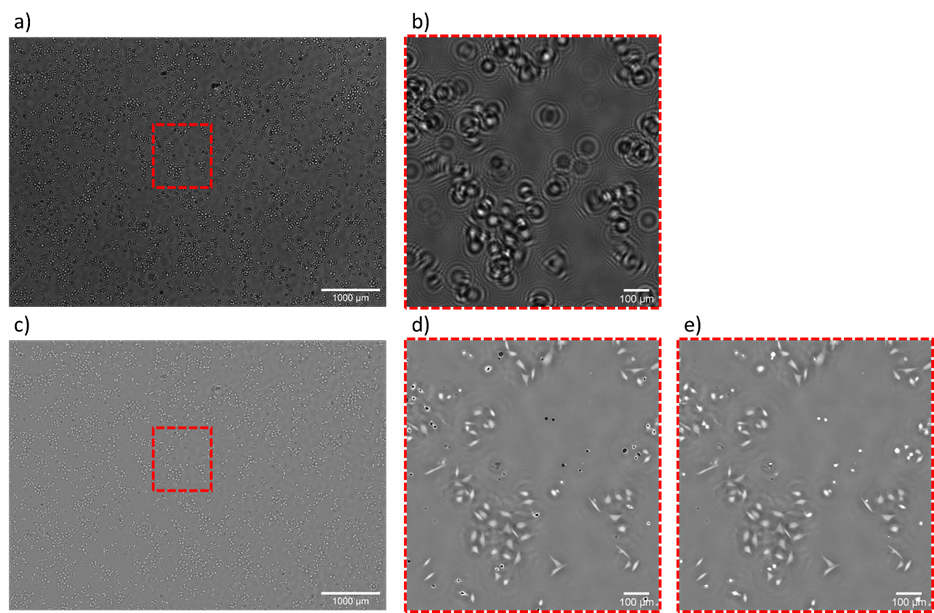

**Supplementary Fig. S2. Holographic reconstruction process**. (a) Raw acquisition of HeLa cells in culture by means of lens-free microscopy (full field of view of 29.4 mm^2^). (b) Detail of (a). (c) Reconstructed phase image (full field of view of 29.4 mm^2^). (d) Detail of (c). (e) Phase image obtained after phase unwrapping. To be compared with (d), before unwrapping.

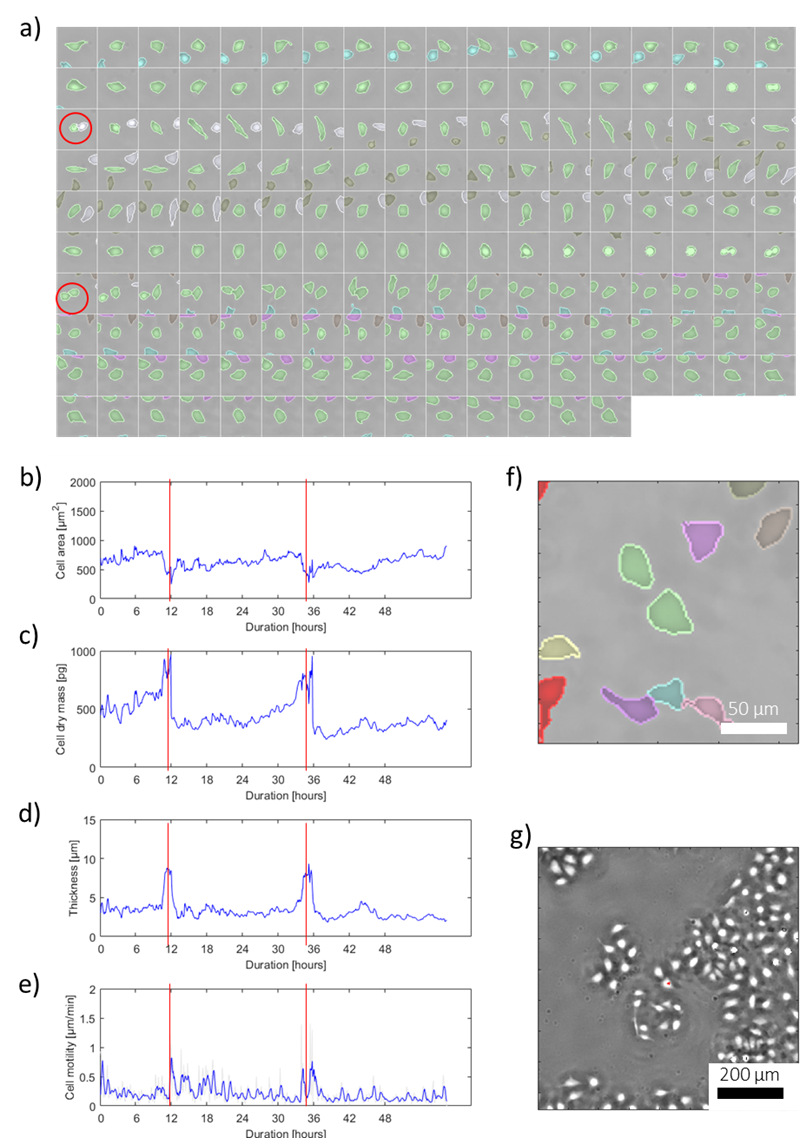

**Supplementary Fig. S3.** Data analysis of a 2.4-days cell trajectory (5 min. frame interval, 710 frames). (a) Cell segmentation performed on the time-lapse acquisition as a function of time (20 minutes interval). The cell of interest is shown with a green overlay. Each cropped image is 75x75 µm^2^. The two cell divisions are shown with red circles. (b) Plot of the cell area as a function of time. (c) Time evolution of the cell dry mass calculated from the integral of the phase over the segmented cell area according to Eqs. (5-7). (d) Plot of the average cell thickness calculated as a function of time. The cell thickness exhibits sharp peaks corresponding to cell divisions (red circles in (a) and red lines in (b-c-d-e)). (e) Plot of the cell motility as a function of time. (f) Segmentation results corresponding to the last frame of the track. The image is 200x200 µm^2^ centered on the cell of interest. (g) 800x800 µm^2^ cropped reconstructed phase image of the last frame. The cell of interest is in the middle with a red spot.

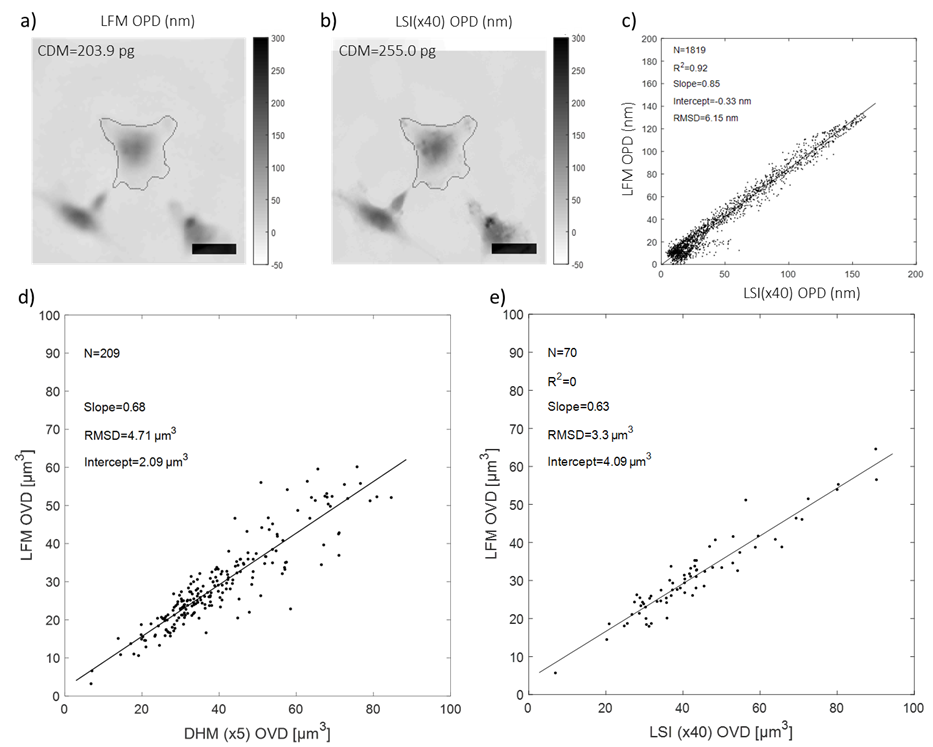

**Supplementary Fig. S4. Cell dry mass measurement.** (a) OPD map (nm) of a fixed COS-7 cell obtained with lens-free microscopy (LFM). Scale bar is 25 µm. The contour of the cell segmentation area is shown in black. (b) OPD (optical path difference) map obtained for comparison with quadriwave shearing lateral interferometry (magnification x40, wavefront sensor SID4Bio Phasics, Saint-Aubin, France). (c) Pixel to pixel comparison between the OPD maps (within segmented cell area). The results of the linear regressions are indicated with values of slope, intercept, coefficient of determination (R^2^) and root-mean-square deviation (RMSD). (d) and (e) Pair-wise comparisons of OVD (see Eq. 6) measurements of fixed COS-7 cells. (d) Comparison between lens-free microscopy and digital holography (magnification x5, DHM T-10105 Lyncée Tec, Lausanne, Switzerland). (e) Comparison between lens-free microscopy and quadriwave shearing lateral interferometry. The linear regression fitting curves are plotted in black. The results of the linear regressions are indicated with values of slope, intercept, coefficient of determination (R^2^) and root-mean-square deviation (RMSD). N refers to the number of cell measurements per comparison.

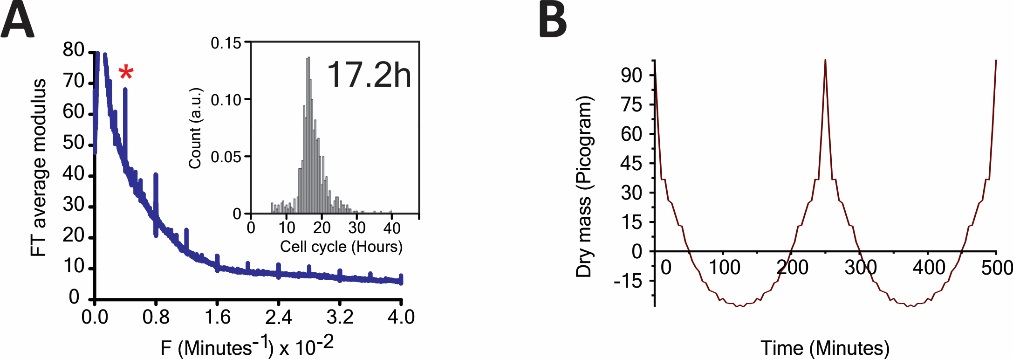

**Supplementary Fig. S5.** (A) The average FT (over 2400 cells) for CHO-K1 cells at 34°C (left) (in the insert, the distribution of the cell cycle times is shown, and mean value is in hr) and (B) the dry mass in units of pg, with effects of noise filtered by inverse FT (right).

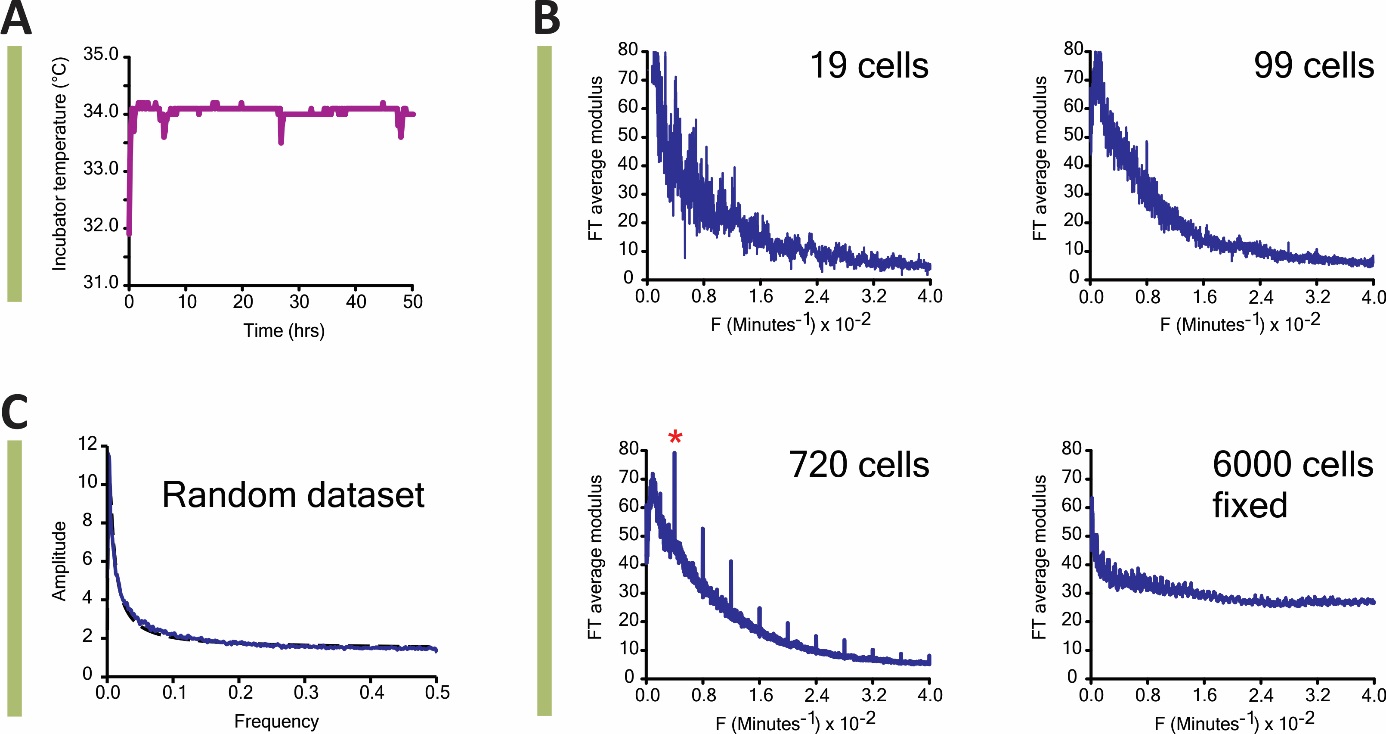

**Supplementary Fig. S6.** (A) The incubator temperature as a function of time; no periodicity is seen. (B) Left, right and lower left: Average FT for increasing numbers of cell trajectories: periodicity appears only as the number of cells increases beyond 100; Lower right: even for 6000 fixed cells there is no rhythm. (C) FT analysis of a random dataset failed to detect any periodicity.

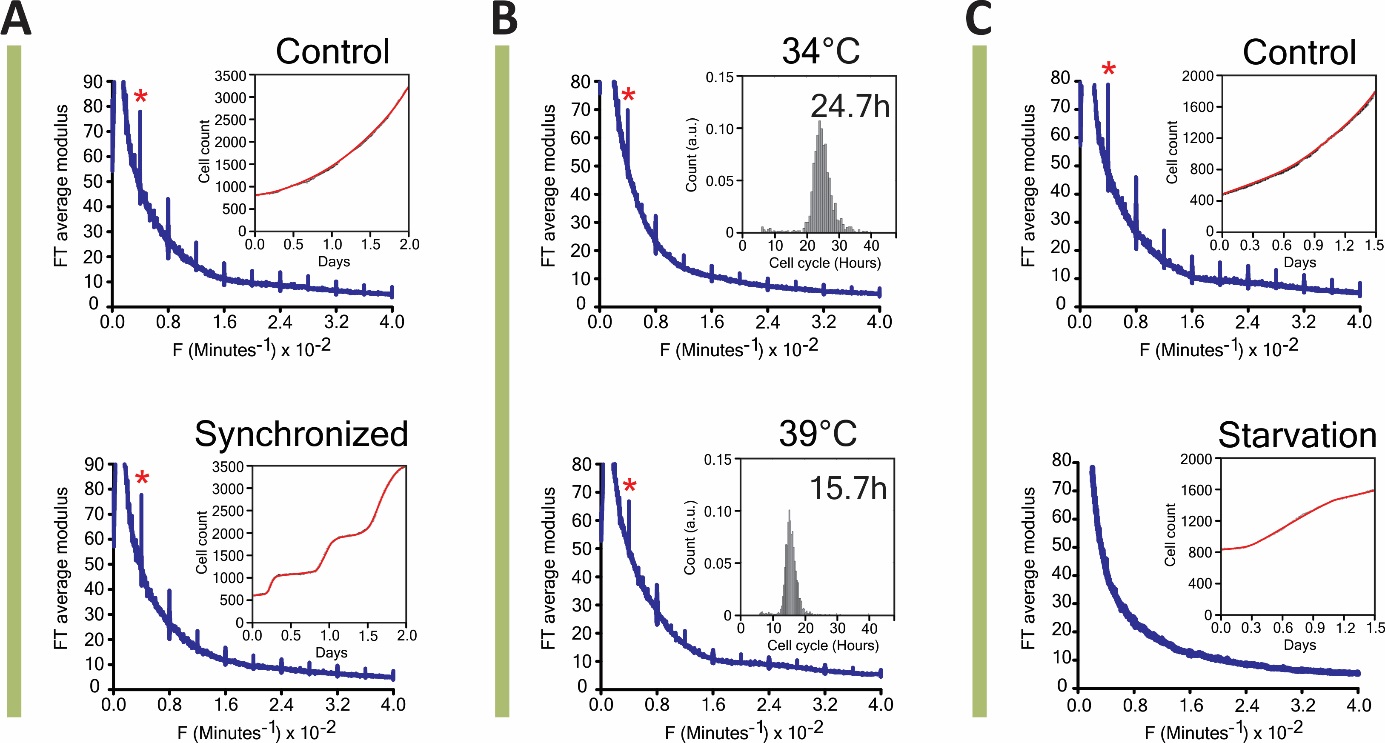

**Supplementary Fig. S7.** (A) Synchronization of HeLa cells using a double thymidine block did not affect the rhythm. In the insert, the cell growth curve is shown for the control and the synchronized cells. The cell growth curve resulted from direct cell count during lens-free observation. (B) Changing the temperature from 34°C (upper) to 39°C (lower) did not change the period although (inserts) the cell cycle lengths were significantly reduced at 39°C (HeLa cells). (C) Serum starvation of Hela cells (lower) eliminates the rhythm compared to control (upper). In insert, the cell count verifies starvation.

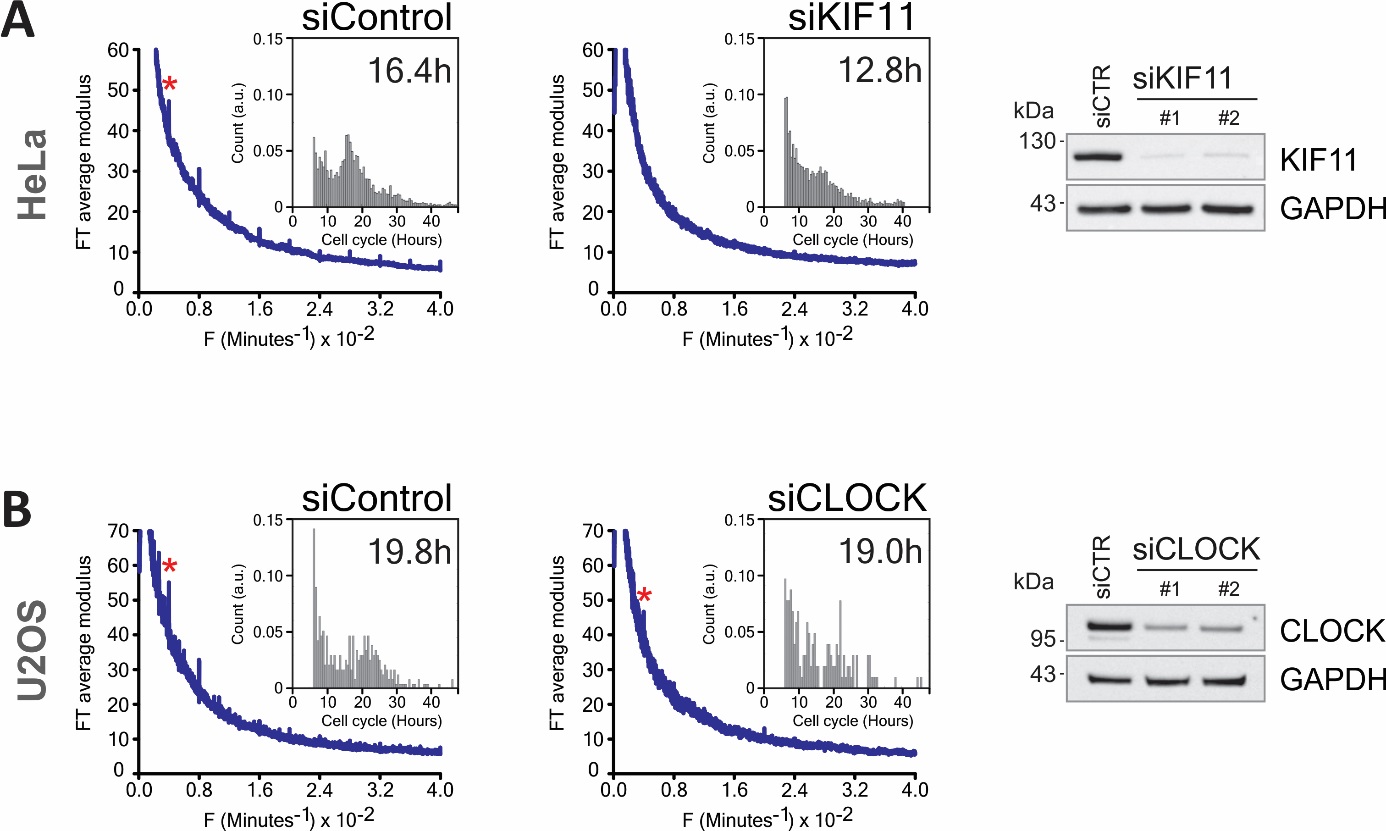

**Supplementary Fig. S8.** (A) When HeLa cells are arrested in G2/M, the rhythm is suppressed. (see insert for the cell cycle length). Western Blots confirms knockdown. (B) Knocking down CLOCK did not eliminate the rhythm in U2OS. In insert, the cell cycle lengths and Western Blot confirm knockdown.

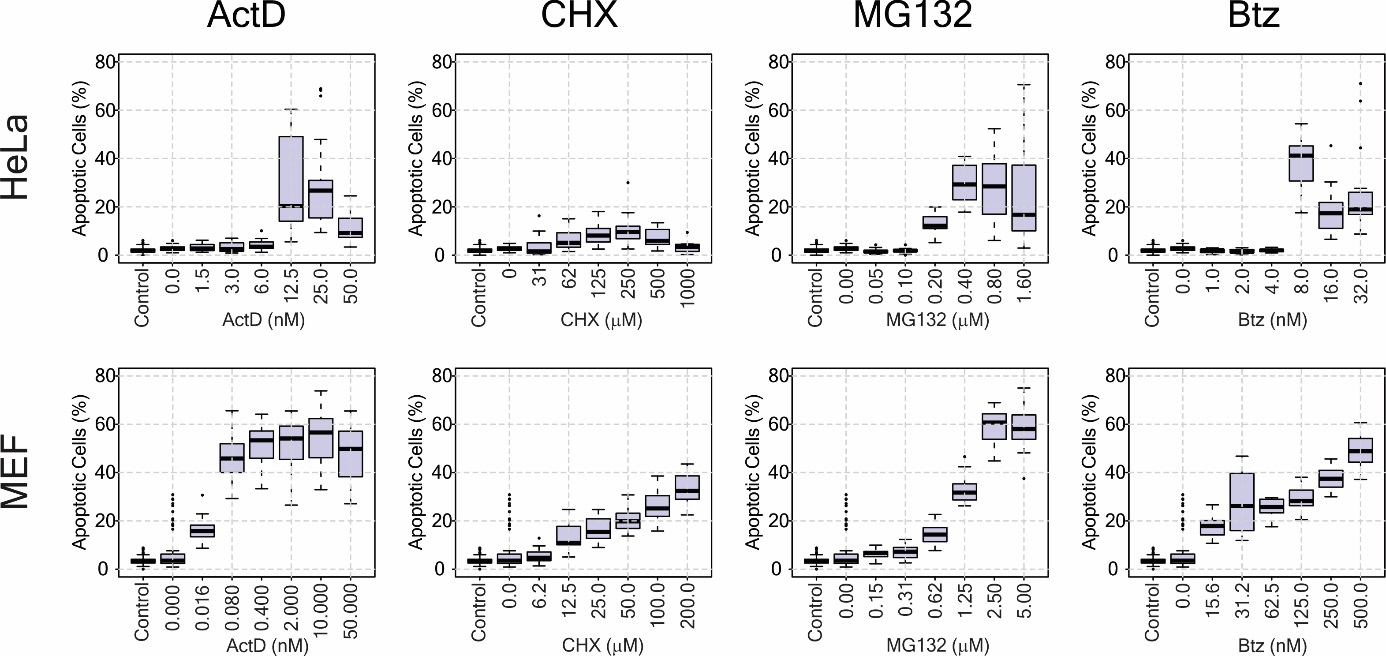

**Supplementary Fig. S9**. Dose response curves for Actinomycin, Cycloheximide, MG132 and Bortezomib in HeLa (top row) and MEF cells (bottom row).

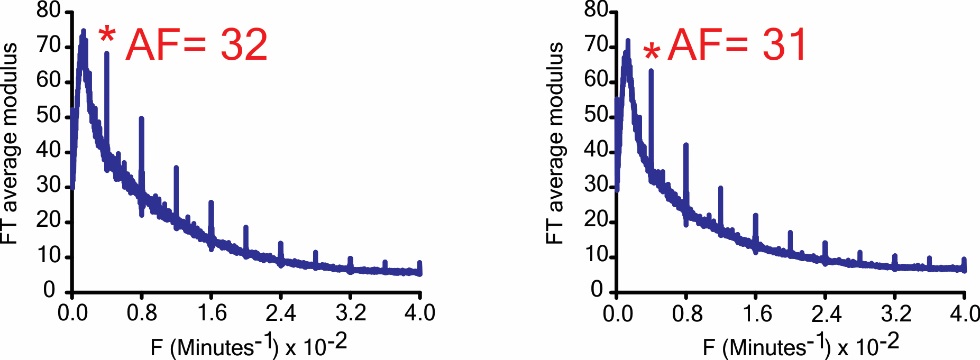

**Supplementary Fig. S10.** Two independent experiments with CHO-K1 cells show the reproducibility of the measurements.

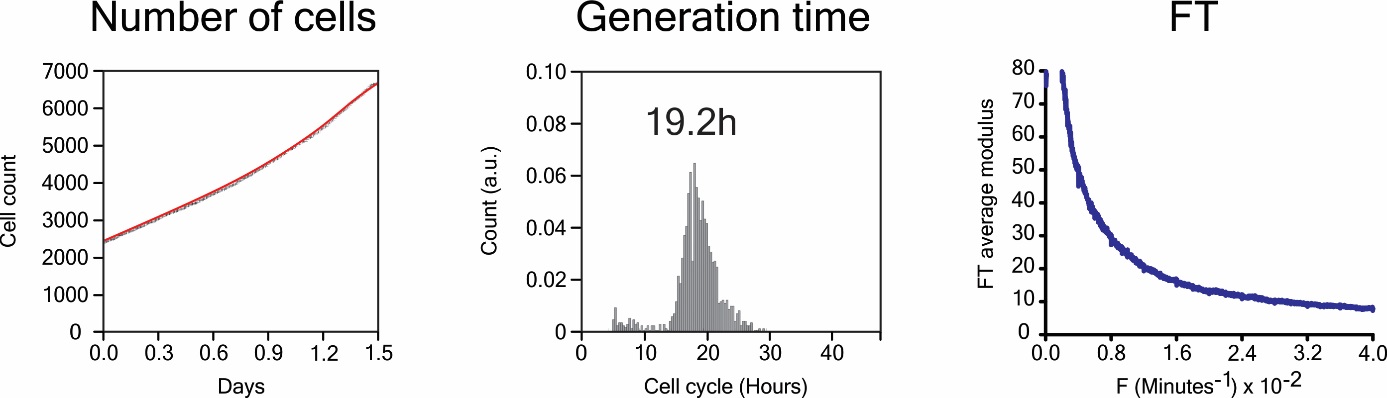

**Supplementary Fig. S11.** HeLa cells treated with 5nM Bortezomib are still cycling, as seen by the increase in cell number (left), with mean cell cycle time estimated as 19.2 hours (middle), but the rhythm is suppressed, as seen by the absence of peaks in the FT (right ).
